# Beyond Student’s t: A Systematic Exploration of Heavy-Tailed Residual Densities for Outlier Handling in Population PK Modeling

**DOI:** 10.64898/2026.03.01.708825

**Authors:** Yiming Cheng, Yan Li

## Abstract

**Background:** Reliable population pharmacokinetic (PopPK) parameter estimation can be compromised by outliers under Gaussian residual error models. A common mitigation strategy is post hoc filtering based on conditional weighted residuals (CWRES); however, this approach can be insensitive due to model “masking” driven by variance inflation. Practical barriers to implementing robust likelihoods in standard software have motivated interest in computationally simpler exponential-tail alternatives such as the Laplace and exponential power distribution (EPD).

**Methods:** We implemented a one-compartment PopPK model using a custom likelihood workaround in Monolix to benchmark four residual error distributions: Normal, Laplace, Generalized Error Distribution (GED), and Student’s t. We assessed CWRES sensitivity under extreme contamination and compared estimation performance using theoretical tail-behavior analysis, controlled simulation studies spanning multiple contamination severities, and a real-world caffeine PK case study with influential terminal-phase deviations.

**Results:** Simulations showed that CWRES-based diagnostics can be unreliable: extreme outliers frequently produced |CWRES| < 6 because the Normal model inflated residual variance, thereby masking contamination. Exponential-tail models (Laplace, GED) improved robustness for mild to moderate outliers but failed under extreme deviations due to insufficiently heavy tails compared to power-law decay. In contrast, the Student’s t model, via power-law tail behavior, maintained stable and minimally biased structural parameter estimates across contamination scenarios. Consistent patterns were observed in the caffeine case study, where the Student’s t model provided improved fit and physiologically plausible parameter estimates.

**Conclusions:** CWRES-driven outlier handling is methodologically fragile because influential contamination can be masked by variance inflation and induce biased inference. Among robust residual error models, exponential-tail distributions may be insufficient for extreme outliers, whereas the Student’s t distribution provides more stable inference across contamination severities. These findings support adopting Student’s t residual modeling as a default robust option in routine PopPK workflows when outlier contamination is plausible.

## Introduction

In pharmacometrics, the reliability and interpretability of population pharmacokinetic (PopPK) parameter estimates depend critically on assumptions about residual variability. ^1^ Standard likelihood-based PopPK workflows most commonly assume that residual errors, the discrepancies between observed concentrations and model predictions, follow a Normal (Gaussian) distribution. ^1, 2^ Although computationally convenient, Gaussian residual models are light-tailed, so large deviations are assigned extremely low probability. ^3^ Consequently, when real-world PK datasets contain outliers, arising from assay variability, protocol deviations (e.g., missed doses or dietary non-compliance), sample handling issues, or transcription/data-entry errors, the Gaussian likelihood can over-penalize those observations, ^4, 5^ This issue is particularly relevant for complex and irregular clinical datasets such as cell and gene therapies, which often exhibit substantially greater variability and a higher frequency of extreme values (apparent outliers) than the conventional PK profiles typically observed with small molecules or monoclonal antibodies. ^6^ During estimation, this can trigger compensatory changes in the model, including drift in clinically interpretable fixed effects (e.g., clearance and volume) and inflation of variability components, biasing inference and degrading interpretability. ^6–8^

Historically, many PopPK analyses have operationalized outlier handling through residual-based diagnostics, most commonly conditional weighted residuals (CWRES), ^9^ combined with heuristic cutoffs (e.g., |CWRES| > 5 or 6) to flag observations for exclusion or sensitivity analyses. ^10–17^ Despite widespread use, this approach has two key limitations. First, the cutoff itself has limited statistical grounding in mixed-effects settings and is often so conservative that only the most extreme deviations would be flagged under idealized assumptions. Second, residual screening can fail when outliers are most consequential: influential observations can induce parameter drift and variance inflation that shrink standardized residuals, masking contamination and rendering CWRES thresholds unreliable as a primary safeguard.

A more principled strategy is likelihood-based robust modeling. Heavy-tailed residual error models provide robustness by assigning non-negligible likelihood to large residuals, reducing the leverage of outlying observations on parameter estimation. ^2, 18–20^ Among these, the Student’s t distribution is widely viewed as a robust default because it is adaptive, making it attractive when the extent of contamination is uncertain. ^6–8, 21^ However, Student’s t residual modeling is not uniformly adopted in applied pharmacometrics, in part may be due to perceived implementation and computational burdens, especially in workflows requiring custom likelihoods, simulation-based evaluation, or sampling-based Bayesian inference. These barriers have motivated interest in simpler heavy-tailed alternatives with closed-form likelihood expressions, such as the Laplace (double-exponential) distribution and the generalized error distribution (GED; exponential-power family). Both are heavier-tailed than Gaussian errors while retaining relatively simple analytic forms, making them appealing in constrained or efficiency-sensitive settings. ^2, 18, 19^

This study addresses a practical methodological question for routine PopPK inference: can simpler exponential-tail residual models such as Laplace or GED provide outlier robustness comparable to Student’s t while reducing implementation burden, or is power-law tail behavior required for stability under clinically realistic contamination? We benchmark Normal, Laplace, GED, and Student’s t residual likelihoods using (i) theoretical density comparisons, (ii) controlled simulations spanning contamination severity, and (iii) a real-world case study using a caffeine PK dataset with influential terminal-phase deviations.

## Methods

### Software and implementation strategy

Data preparation and post-processing were performed in RStudio (R Version 4.1.3, Posit Software, PBC, Boston, MA, USA), and all model fitting was conducted in Monolix® (Version 2023R1, Lixoft SAS, Antony, France). A key technical limitation of Monolix is that user-defined residual-error likelihoods are not supported for standard continuous outcomes; custom likelihood specification is only available for integer-valued count/categorical models. To bypass this restriction, we used the count datatype solely as an implementation vehicle while preserving a continuous-data likelihood formulation. Specifically, observed concentrations were multiplied by 10^6^ and rounded to integers prior to import so that Monolix would accept the dependent variable as a count outcome. Within the Monolix model code, the integer-valued observations were immediately mapped back to the original continuous scale by dividing by10^6^. The residual-error likelihood (Normal, Laplace, GED, or Student’s t) was then evaluated on this rescaled continuous quantity, enabling estimation under the intended continuous probability density function while leveraging the count-model interface required to implement a custom likelihood in Monolix. The 10^6^ scaling factor was chosen to retain sufficient numerical precision during integer encoding.

Monolix (rather than NONMEM) was used to reflect a practical implementation constraint: whereas NONMEM readily supports a broad range of user-defined likelihood formulations for continuous outcomes, Monolix imposes restrictions that make implementation of robust residual models non-trivial. This setting therefore provides a relevant test bed for evaluating whether computationally simple heavy-tailed residual distributions with closed-form pdf/cdf can be deployed with minimal engineering overhead in software environments without native support for robust likelihoods.

### Model structure and simulation design (Data Simulation)

Simulated datasets were generated and subsequently fitted using four residual-error likelihoods to evaluate performance under controlled outlier contamination. One-compartment oral pharmacokinetic (PK) profiles were simulated for 50 virtual subjects following a single 400-mg oral dose. Structural parameters were fixed to an absorption rate constant ka = 0.25 hr^−1^, apparent clearance CL/F = 5.3 L/hr and apparent central volume V/F = 36 L. Interindividual variability (IIV) was assumed log-normal for all structural parameters with 20% coefficient of variation (i.e., ω = 0.2 on the log scale for each parameter). Residual unexplained variability was modeled using a proportional error model with l1 = 0.2.

To generate outlier-contaminated profiles, a single terminal-phase concentration observation was perturbed by multiplying the observed value by a prespecified factor. A factor of 1 was used for the baseline simulation (no outliers). For the outlier-contamination simulations, multiplicative factors of 5–20 were used to represent moderate deviations, and 30–100 to represent extreme deviations. This design preserves the underlying PK trajectory while introducing controlled, phase-specific contamination intended to mimic influential terminal observations that can drive apparent clearance misestimation.

Each simulated dataset was fitted under four residual-error likelihood specifications: Gaussian (Normal), Laplace, GED, and Student’s t. Model performance was compared across contamination severities based on estimation accuracy at both the population and individual parameter levels, complemented by visual inspection of individual PK profiles with observed concentrations overlaid against model predictions.

### Residual-error likelihoods evaluated

Four residual-error probability density functions (PDFs) were evaluated. The Normal (Gaussian) model (Eq. 1) corresponds to a squared-error loss and therefore penalizes large deviations quadratically. ^22^ The Laplace (double exponential) model (Eq. 2) corresponds to an L1-norm loss and down-weights large deviations relative to the Gaussian. ^22^ The GED model (Eq. 3) introduces a shape parameter n that controls tail thickness; smaller n yields heavier tails, with n = 2 reducing to the Gaussian and n = 1 reducing to the Laplace. ^23^ The Student’s t model (Eq. 4) estimates the degrees of freedom ν and exhibits power-law tail decay, providing strong robustness to extreme deviations as ν decreases. ^21^

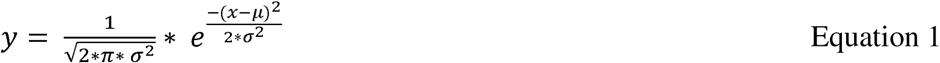

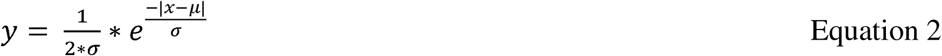

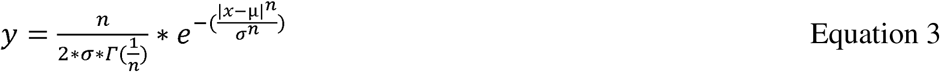

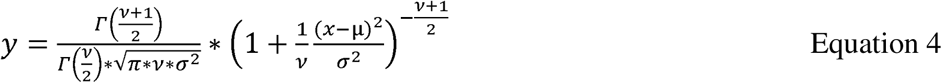

where x denotes the observed value, μ denotes model-predicted value, n denotes the GED shape parameter, and ν denotes the degrees of freedom for Student’s t distribution. Г is the gamma function.

### Real-world case study

To corroborate the simulation findings with real-world data, we analyzed a caffeine PK dataset from a clinical drug–drug interaction study in patients with acute myeloid leukemia (AML) and myelodysplastic syndrome (MDS). ^7^ The dataset included 19 subjects who received a single 100-mg oral dose of caffeine under two conditions: in the absence of the perpetrator drug (Period 1) and in its presence (Period 2). In Period 1, several subjects exhibited unexpectedly high caffeine concentrations at 24 and/or 30 hours post-dose for reasons that could not be definitively determined. Under conventional analyses, such terminal-phase observations would typically be treated as outliers and, if not handled appropriately, could materially influence the assessment of the perpetrator effect on caffeine exposure.

In prior internal work, we applied a Student’s t residual-error model to improve robustness of PopPK inference in this setting. In the present study, the same dataset was re-fitted using four residual-error likelihood specifications: Normal, Laplace, GED, and Student’s t. Model performance was evaluated by comparing observed and model-predicted concentration–time profiles at the individual-subject level, with emphasis on subjects exhibiting influential terminal-phase deviations. This case study was used to assess, in an applied setting, the relative ability of each residual-error likelihood to accommodate real-world outliers without destabilizing parameter estimation or distorting model-based predictions.

## Results

### Impact of Terminal-Phase Outliers on CWRES and PK Parameter Estimation

Figure 1 demonstrates that standard residual-based outlier screening using conditional weighted residuals (CWRES) can fail to identify highly influential terminal-phase outliers in PopPK analyses. Figure 1A shows concentration–time profiles for five representative simulated subjects in which a single observation at 48 h (terminal phase) was artificially inflated by a 20-fold multiplicative factor, an intentional outlier contamination designed to exert strong leverage on the terminal slope and, therefore, clearance. Despite the obvious deviation in the raw profiles, the corresponding CWRES time courses (Figure 1B) did not flag these observations under commonly used criteria. In our examples, the CWRES values at 48 h remained well below 6 and were often <3. Under typical pharmacometric practice, these points would therefore be retained and implicitly treated as consistent with the assumed residual error model. The consequence of retaining such terminal spikes is evident in the individual fits (Figure 1C): to accommodate the inflated 48 h concentrations, the fitted elimination phase (red curves) bends upward relative to the true trajectory (green curves), flattening the terminal slope and systematically biasing clearance downward. This distortion also propagates to the population level. As summarized in Table 1, both ka and V were substantially overestimated (0.50 vs 0.25; 68.4 vs 36.0), clearance was underestimated (3.96 vs 5.30), and the residual error was markedly inflated (0.104 vs 0.040), consistent with the model absorbing the outlier impact through parameter drift and variance inflation rather than isolating the contaminated point. To assess how severe contamination must be before CWRES becomes informative, we repeated the experiment across a series of multiplicative factors (e.g., 5, 30, 50, 75, and 100; Supplementary Figure S1). Across this range , including the most extreme outlier contamination, CWRES remained below typical cutoffs while both individual- and population-level estimates deviated materially from the known truth, indicating that standardized residual thresholds can be an unreliable safeguard when influential outliers induce poor model performance such as variance inflation and structural parameter drift.

**Figure 1.**
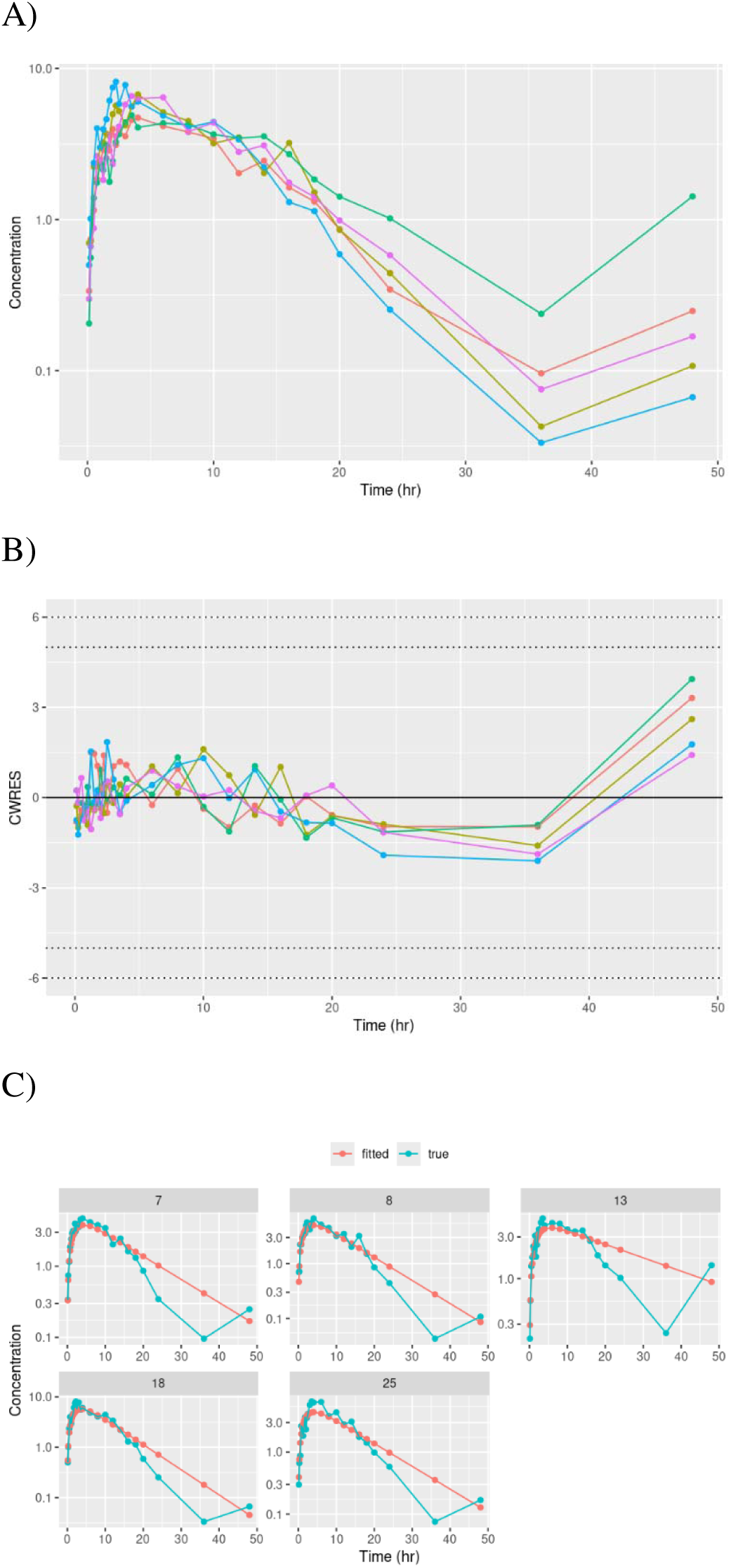
Impact of influential terminal-phase outliers on CWRES diagnostics and pharmacokinetic modeling. (A) Simulated concentration–time profiles for five representative subjects. (B) Corresponding CWRES time courses for the same subjects. (C) Observed concentration–time profiles (green) overlaid with model-predicted profiles (red) for the same subjects. Terminal-phase observations were inflated by a 20-fold multiplicative factor. Dotted horizontal lines denote the |CWRES| thresholds of 5 and 6.

**Table 1.** Population PK parameter estimates under terminal-phase outlier contamination across multiplicative inflation factors, illustrating structural parameter drift and residual-variance inflation under the Gaussian residual model.

| PK parameter | True Value | Multiplicative Factor |  |  |  |  |
| --- | --- | --- | --- | --- | --- | --- |
|  |  | 5 | 30 | 50 | 75 | 100 |
| CL/F (L/hr) | 5.30 | 5.09 | 3.70 | 2.74 | 2.00 | 1.41 |
| ka (1/hr) | 0.25 | 0.40 | 0.55 | 0.60 | 0.64 | 0.64 |
| V/F (L) | 36.00 | 53.05 | 74.46 | 78.72 | 86.11 | 84.67 |
| $\sigma^2$ | 0.04 | 0.07 | 0.12 | 0.15 | 0.17 | 0.19 |

Multiplicative factor denotes the fold-inflation applied to a single terminal-phase concentration observation per dataset. Values are population parameter estimates from the Gaussian residual-error model for each contamination scenario.
Abbreviations: CL/F, apparent clearance; V/F, apparent central compartment volume; $\sigma^2$ , variance of the proportional residual error.

### Tail Behavior of Exponential-Tail Versus Power-Law Residual Models

Figure 2 compares the probability density functions of the Laplace and generalized error (GED/exponential-power) families across a range of shape parameters n, overlaid with the Normal distribution and Student’s t distributions with df = 2 and 5. These candidate residual models differ primarily in how rapidly the likelihood decays as the standardized residual magnitude | z | increases, which determines the penalty assigned to extreme deviations during estimation, where 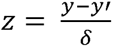. Under the Gaussian model, the log-density is quadratic in *z*, leading to rapid tail decay (*f*(*z*) ∝ exp (-*z*^2^/2)). In contrast, the Laplace and GED families exhibit exponential-tail decay, with log-density increasing approximately linearly or sublinearly in |z| depending on the shape parameter (*f*(*z*) ∝ exp (-| z |^n^)). Within this family, smaller n produces heavier tails: n = 1 corresponds to the Laplace (double-exponential) distribution, whereas n < 1 yields heavier tails than Laplace, consistent with the visibly higher tail densities in Figure 2. By comparison, the Student’s t distribution exhibits power-law behavior; for large |z|, the density decreases polynomially (*f*(*z*) ∝ | z |^−(v+1))^, leading to substantially greater tail mass than exponential-tail models at extreme residual magnitudes.

**Figure 2.**
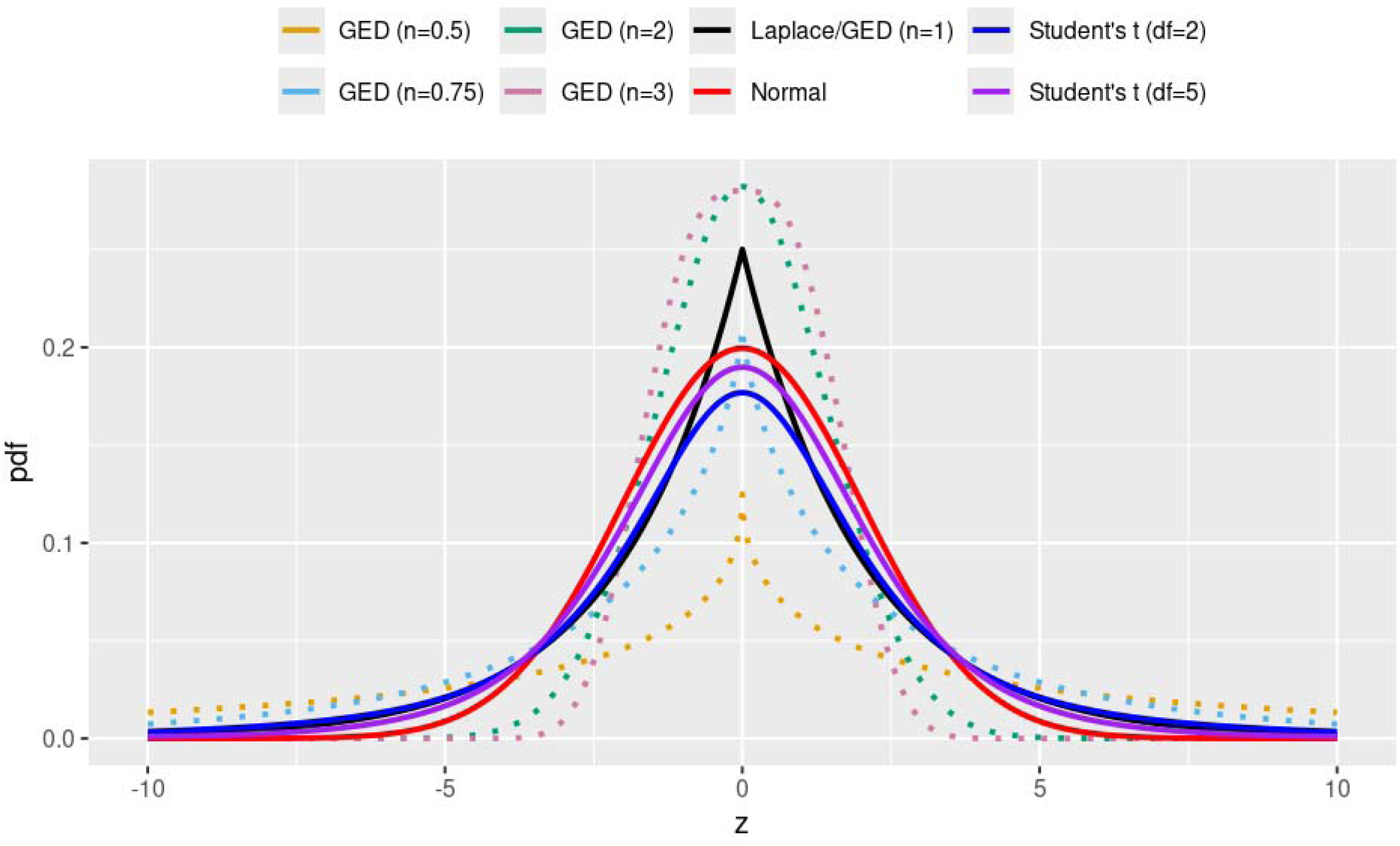
Probability density functions of the Laplace and generalized error (GED/exponential-power) families across shape parameters *n*, overlaid with the Normal distribution and Student’s t distributions (ν = 2 and ν = 5).

Visual inspection of Figure 2 suggests two practically relevant findings. For moderate deviations, Laplace and heavy-tailed GED models (n ≤ 1) assign higher density than the Normal model, indicating weaker penalization of moderate residual excursions. For larger deviations, the Student’s t curves maintain comparatively higher tail density; even with df = 5, Student’s t remains above Laplace/GED as |z| increases, with a more pronounced separation at df = 2. Collectively, Figure 2 highlights meaningful differences in tail decay across candidate residual models. Importantly, Laplace and GED likelihoods retain simple closed-form expressions, whereas Student’s t achieves heavy tails through a different complicated functional form and can be less straightforward to implement in workflows requiring user-defined likelihoods. These observations motivate the subsequent empirical comparisons: we evaluate whether exponential-tail alternatives such as Laplace/GED can offer practical robustness to outliers comparable to Student’s t in PopPK estimation, while potentially reducing implementation burden and computational cost in simulation- and sampling-based workflows.

### Simulation Study: Baseline Performance Without Outliers

Figure 3A summarizes population-level parameter estimates under the baseline “clean-data” simulation setting, with proportional inter-individual variability (IIV) and proportional residual variability each set to 0.2 during dataset generation. Across all four residual error models (Normal, Student’s t, Laplace, and GED), fixed-effect parameters (CL, V and ka) and variance components were recovered consistently and remained close to the true values (red reference lines), indicating negligible bias when the dataset contains no gross contamination. In this setting, the Student’s t model estimated a large degrees-of-freedom value (∼25), which corresponds to near-Gaussian behavior and confirms that the t-likelihood adaptively collapses toward the Normal model when heavy-tailed behavior is not supported by the data. Similarly, the GED shape parameter converged toward values consistent with approximately Gaussian errors, further indicating that the additional flexibility of the robust families did not induce spurious heavy-tailed behavior in clean datasets.

**Figure 3.**
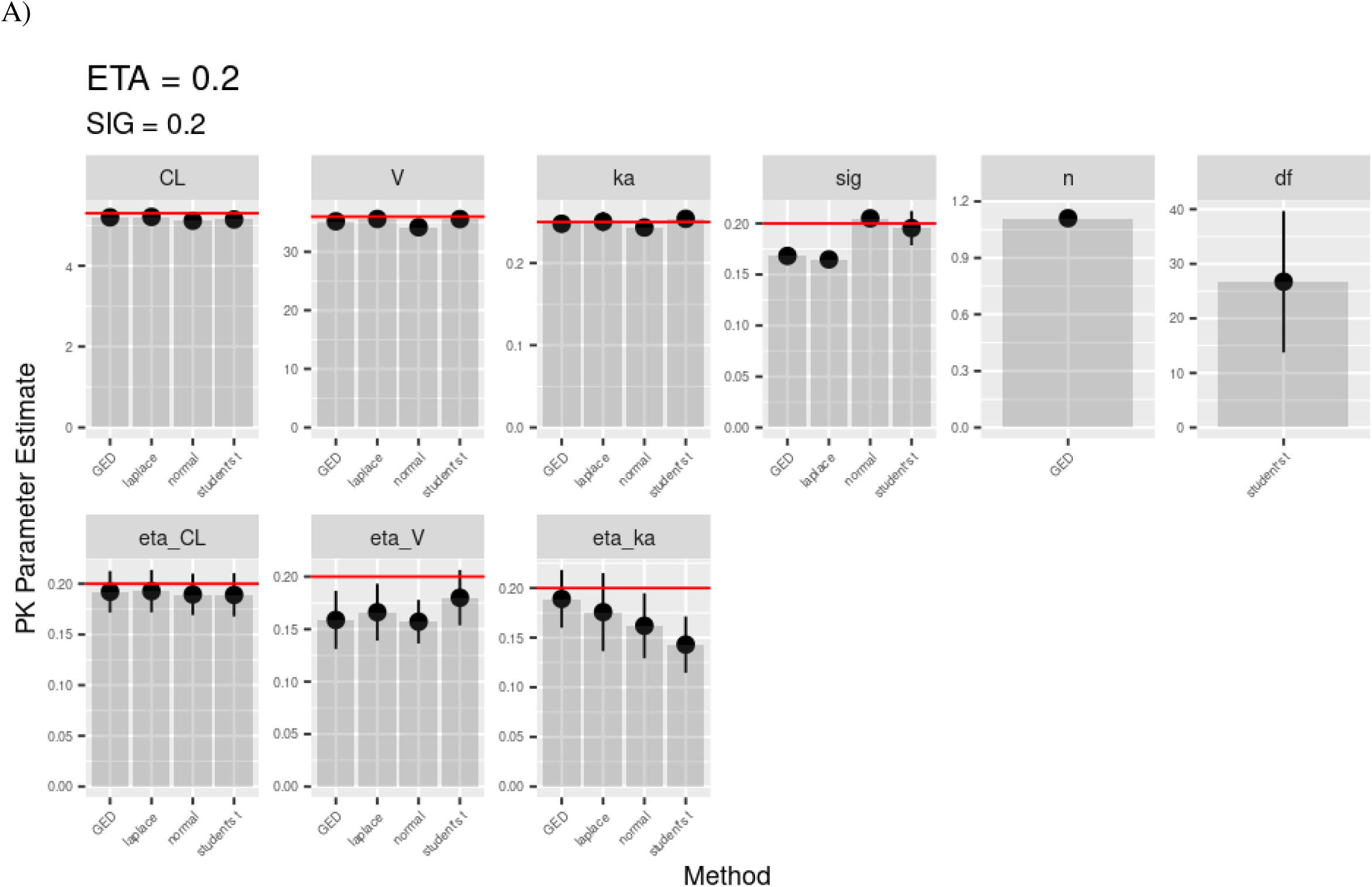
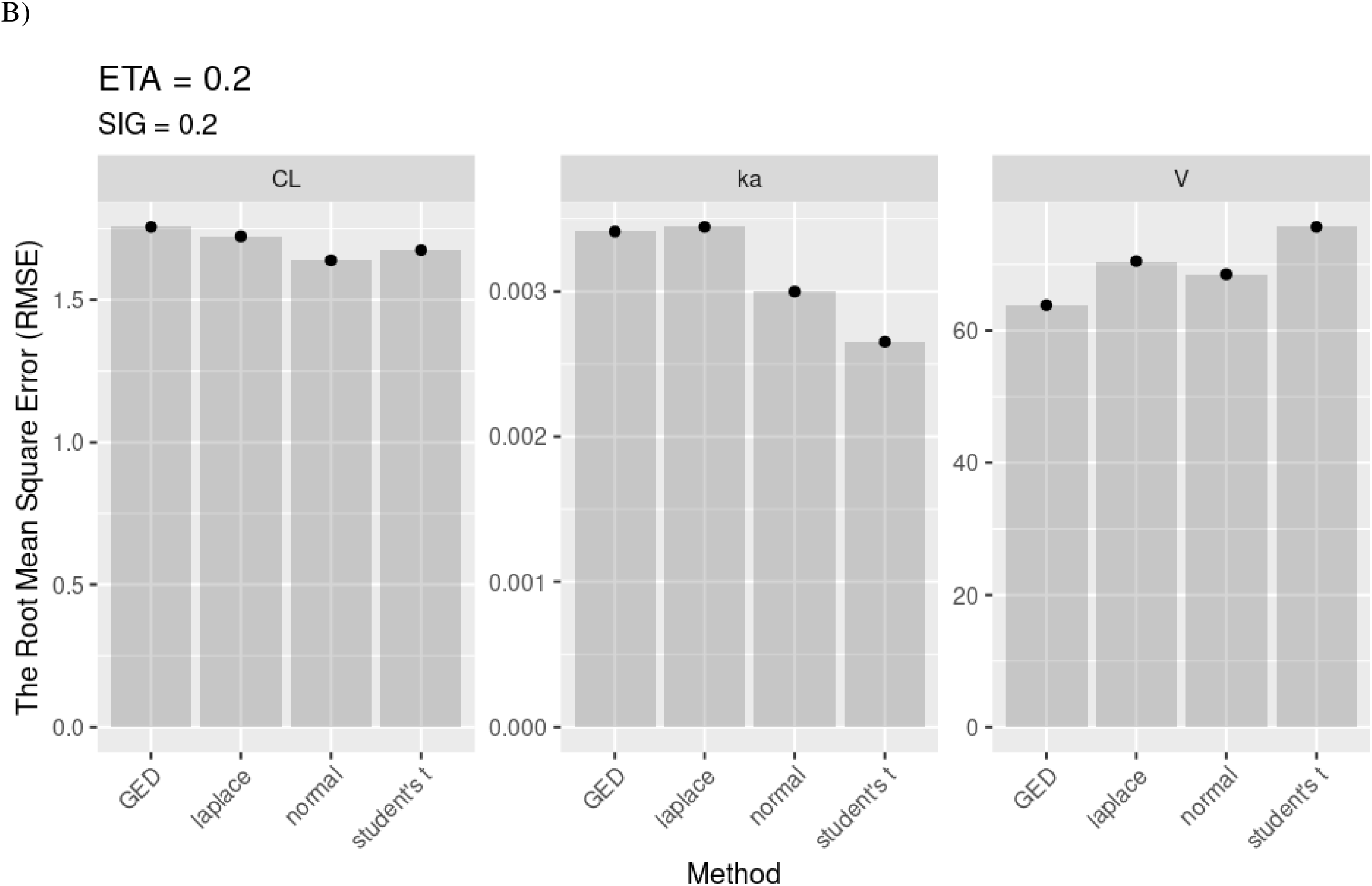
Baseline simulation without outliers: comparable population- and individual-level PK parameter recovery across Normal, Student’s t, Laplace, and GED residual-error models. (A) Population-level parameter estimates under the clean-data setting. (B) Subject-level accuracy summarized by root mean square error (RMSE) of individual parameter estimates relative to the true simulated values across subjects. Datasets were generated with proportional inter-individual variability (IIV) = 0.2 and proportional residual variability = 0.2. Red lines indicate the true parameter values used for simulation.

Figure 3B provides the complementary subject-level assessment using root mean square error (RMSE) of the individual parameter estimates relative to the true simulated values across all subjects. Consistent with the population-level findings, RMSE values for CL, V and ka were comparable across the Normal, Student’s t, Laplace, and GED models, demonstrating that none of the robust residual specifications compromised estimation accuracy in the absence of outliers.

To assess generalizability, we repeated the clean-data evaluation across a wider range of variability settings by combining varying both proportional IIV and proportional residual error (e.g., 0.2, 0.4, and up to 0.6). Across these scenarios (Supplementary Figures S2A–S2B), the same qualitative pattern was observed: all four residual models achieved stable estimation and similar parameter recovery when outliers were absent. Collectively, these results support that introducing robust residual families (Student’s t, Laplace, GED) does not degrade performance in clean PopPK datasets, establishing an appropriate baseline for the subsequent outlier-contamination experiments.

### Simulation Study: Performance Under Outlier Contamination

Figure 4A summarizes population-level parameter estimates under a moderate outlier contamination setting (multiplier = 12). In this scenario, the Normal model showed clear sensitivity to contamination, most notably through inflation of the residual error and associated variance components, consistent with the estimation algorithm attempting to accommodate extreme observations by increasing unexplained variability rather than isolating their influence. In contrast, the heavy-tailed residual models (Laplace, GED/exponential-power, and Student’s t) provided more stable estimates of σ and better controlled the inflation of variability terms. For fixed effects, Student’s t generally achieved the closest recovery of the true PK parameters relative to the exponential-tail alternatives. While Laplace and GED showed modest improvement over the Normal model for *cL*, they did not consistently correct bias in *k_a_* and *v*, and their inter-individual variability (IIV) estimates remained more distorted than those obtained under Student’s t, indicating limited robustness when outliers are both frequent and influential.

**Figure 4.**
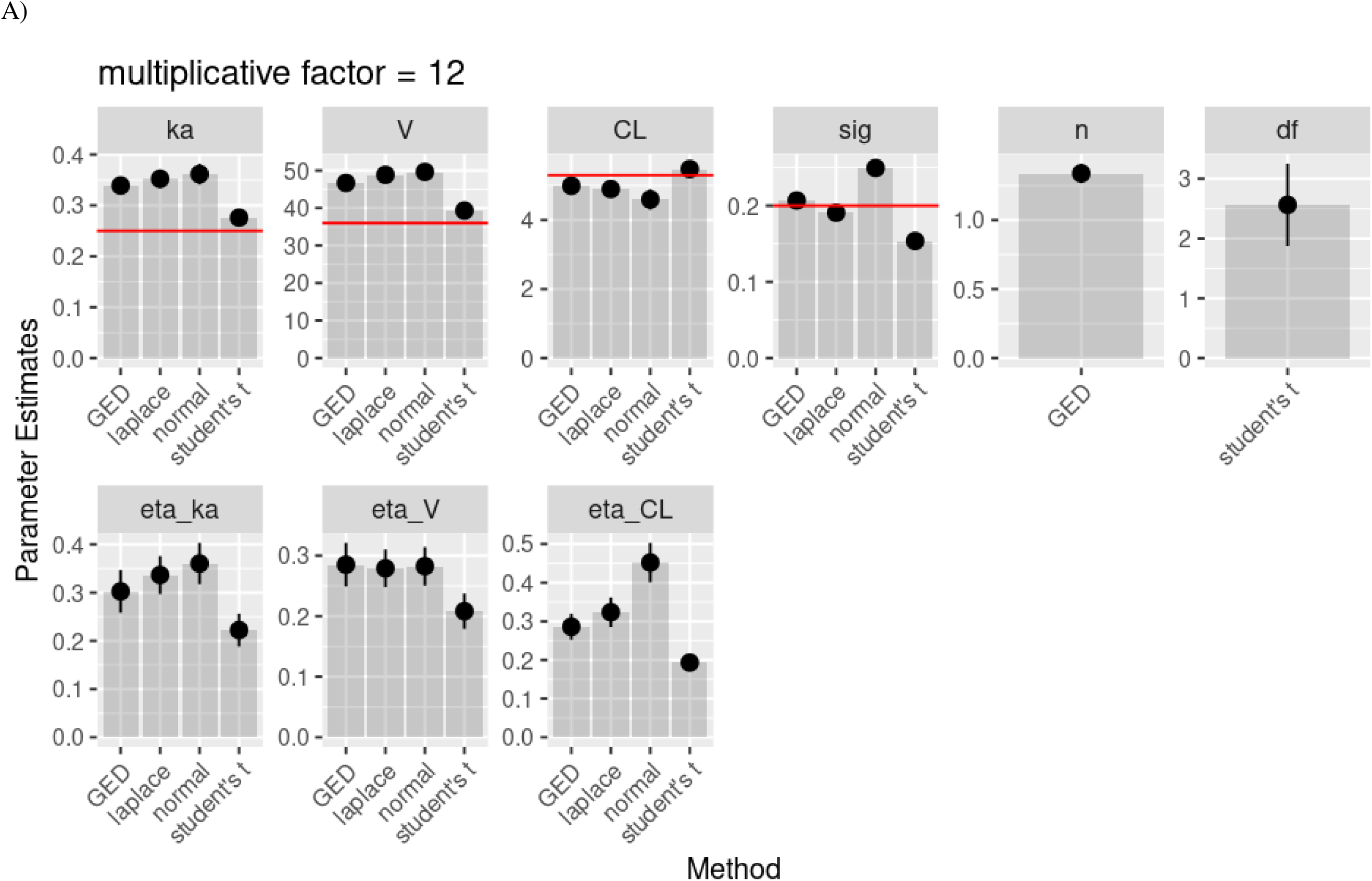
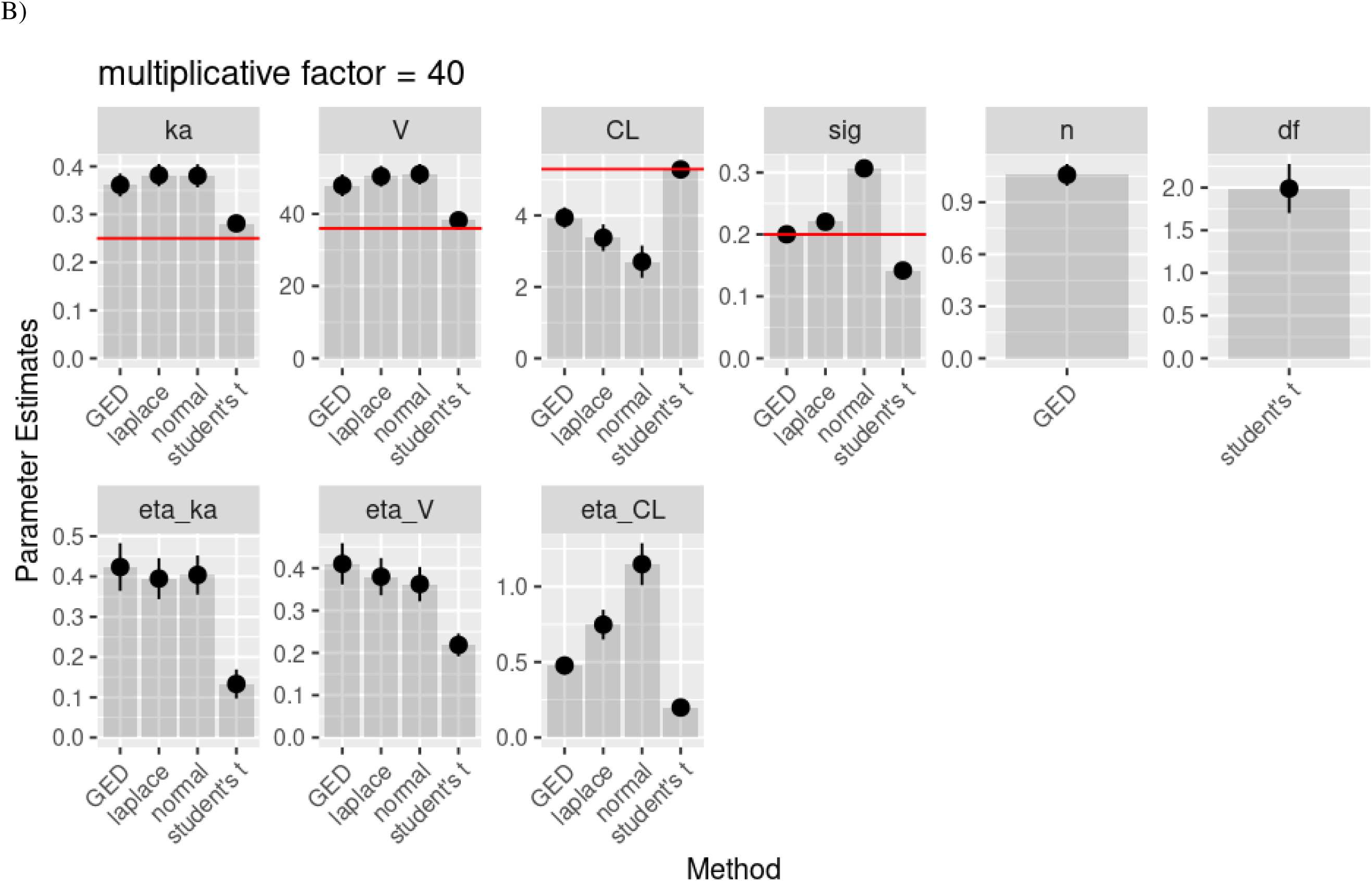
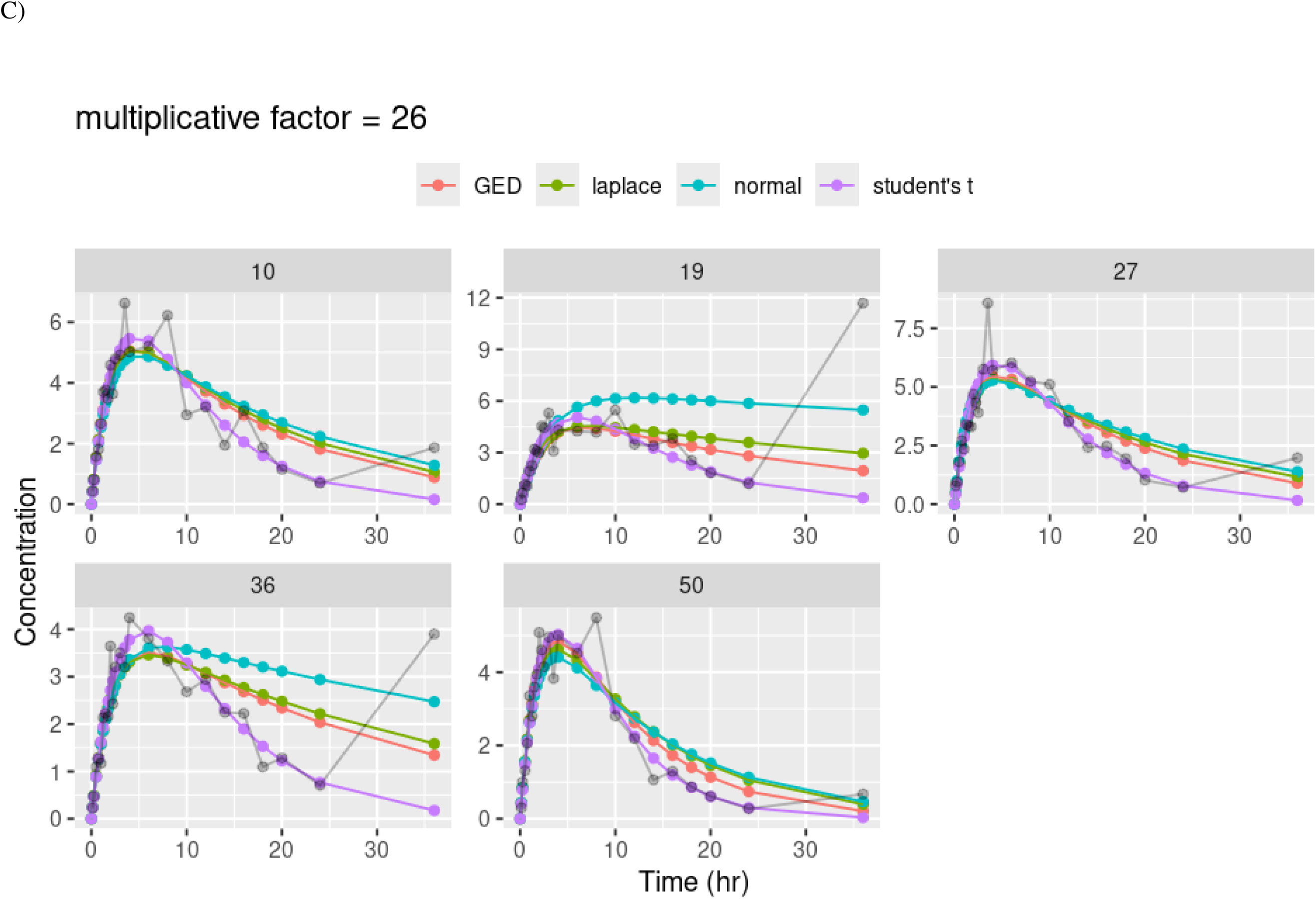
Simulation performance under moderate and severe terminal-phase outlier contamination: Student’s t yields the most stable population parameter recovery and individual fits compared with Normal, Laplace, and GED residual-error models. (A) Population-level parameter estimates under moderate contamination (terminal-phase multiplier = 12). (B) Population-level parameter estimates under severe contamination (terminal-phase multiplier = 40). (C) Representative individual fits under moderate contamination (multiplier = 12): observed concentration–time profile (black) overlaid with model fits from GED (red), Laplace (green), Normal (blue), and Student’s t (purple).

Figure 4B extends the comparison to a more severe contamination setting (multiplier = 40). As expected, the separation among methods became more pronounced. Student’s t maintained the most consistent parameter recovery across fixed effects and variability components, whereas the Normal model exhibited substantial degradation. Laplace and GED continued to provide partial protection relative to the Normal likelihood, most visibly for *CL*, but both exponential-tail models still showed appreciable bias for *k_a_, v*, and several variance components under this stronger outlier contamination, suggesting that exponential-tail heaviness alone may be insufficient to preserve stable estimation under severe contamination. Additional sensitivity analyses across contamination scenarios are provided in Supplementary Figure S3 and show consistent qualitative patterns.

Figure 4C illustrates representative individual fits under the contaminated setting. Student’s t (purple curves) yielded the most stable individual trajectories in the presence of extreme observations. By contrast, under the Normal model, fitted profiles were pulled toward the contaminated late-time points, with the elimination phase bending upward relative to the underlying trajectory (black), effectively flattening the terminal slope and biasing clearance downward. Laplace and GED reduced, but did not fully eliminate, this profile distortion, producing fits that were less influenced by the outliers than the Normal model yet still showing residual susceptibility under the most severe contamination.

### Real-World Case Study: Caffeine PK Dataset From a Drug–Drug Interaction Study

Supplementary Figure S3 displays individual caffeine concentration–time profiles from a real-world drug–drug interaction case study, with Period 1 and Period 2 corresponding to caffeine PK in the absence and presence of the perpetrator, respectively. Several subjects (notably IDs 1, 4, 6, and 16) exhibit anomalously elevated concentrations at the last scheduled time point in Period 1, consistent with influential terminal-phase deviations that are unlikely to be explained by the structural model alone. Building on our prior Student’s t analysis of this dataset, we re-fit the same data using Laplace and GED residual likelihoods to directly compare exponential-tail alternatives with Student’s t, alongside the standard Normal model.

Figure 5 presents representative individual fits under each residual specification. Across the highlighted subjects, the Student’s t model provided the most stable and physiologically plausible characterization of the terminal phase in the presence of the 30 h spikes, avoiding disproportionate distortion of the elimination slope. In contrast, the Normal model was most susceptible to these late-time deviations, with fitted profiles pulled upward toward the extreme observations. The Laplace and GED models demonstrated intermediate behavior: relative to the Normal likelihood, both exponential-tail models reduced the influence of the terminal spikes and improved agreement with the bulk of the data, but they still showed residual sensitivity when deviations were large, and their correction did not match the robustness observed under Student’s t for these subjects.

**Figure 5.**
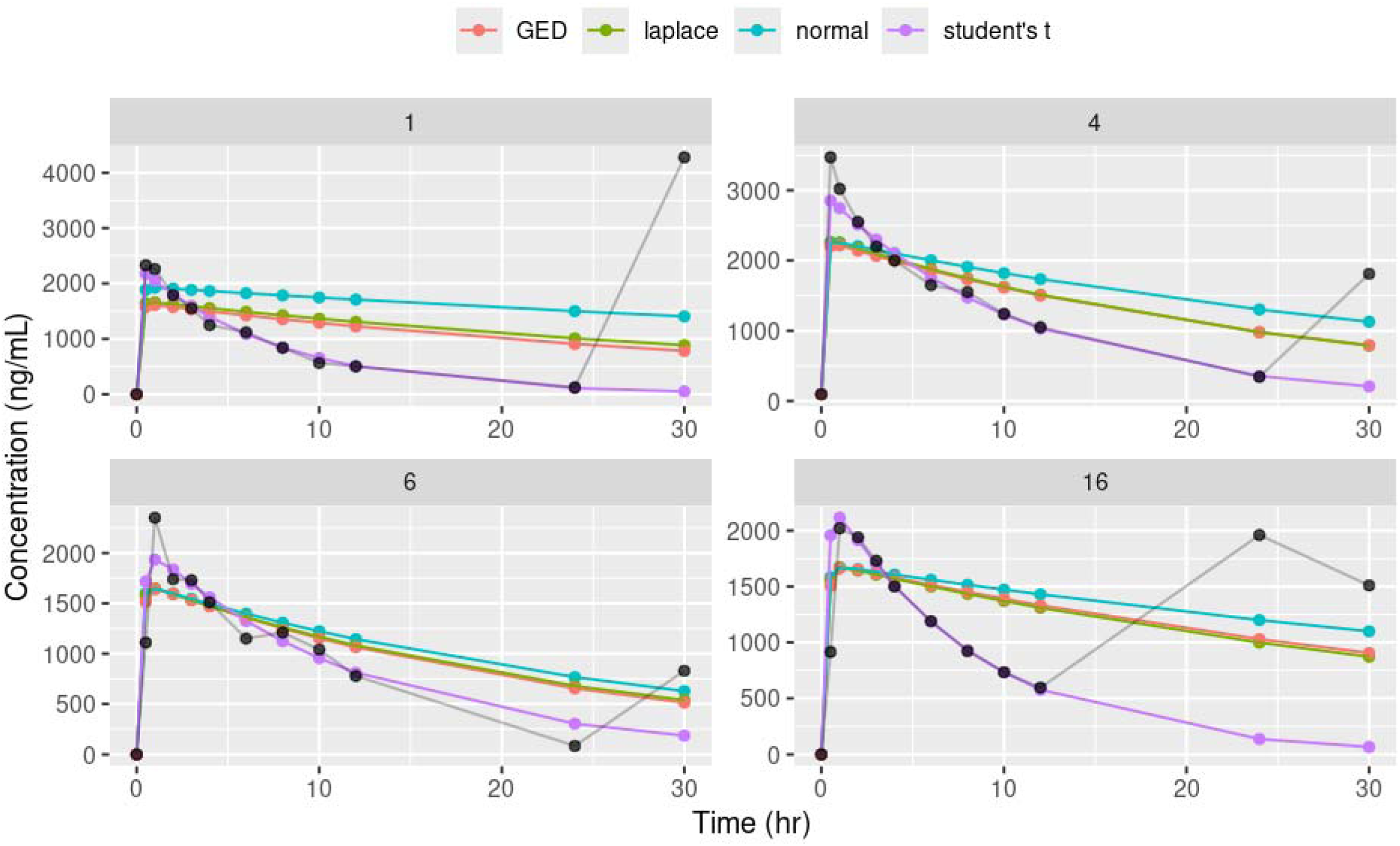
Representative individual caffeine PK fits comparing residual-error models: Student’s t provides the most stable terminal-phase characterization in the presence of 30-h concentration outlier contaminations. Observed concentration–time profile (black) overlaid with model fits from GED (red), Laplace (green), Normal (blue), and Student’s t (purple).

Taken together with the simulation findings, this case study indicates that real clinical PK datasets can contain terminal-phase deviations of sufficient magnitude that exponential-tail likelihoods (Laplace/GED) provide only partial protection, whereas Student’s t more effectively accommodates extreme residual behavior.

## Discussion

Traditional maximum likelihood estimation (MLE) remains the cornerstone of pharmacometric modeling, ^24^ but it is intrinsically vulnerable to the outliers that are unavoidable in real-world pharmacokinetic datasets. ^1, 4^ Such outliers may arise from assay variability, protocol deviations (e.g., missed doses or dietary non-compliance), sample handling issues, or transcription/data-entry errors. Under the default Gaussian residual model commonly used in the pharmacometrics field, the squared-error penalty causes aberrant observations to exert disproportionate leverage on the objective function, which can bias fixed-effect estimates, distort variability components, and inflate uncertainty. ^1^ In routine practice, this sensitivity is often addressed through residual-based screening, most commonly conditional weighted residuals (CWRES), where observations exceeding a heuristic cutoff (often |CWRES| > 6) are flagged for potential exclusion or further assessment. ^3, 9^ This approach is problematic on two levels. First, it lacks a meaningful statistical basis in this setting: if CWRES were truly standard Normal, |CWRES| cutoffs of 5 or 6 would correspond to an exceedingly rare event (two-sided probability approximately 5.7×10^−7^ and 2×10 , respectively), making it so conservative that it can only detect the most extreme deviations. Second, and more importantly, our results show that standardized residual screening is not a dependable safeguard against influential outliers due to model masking. In our simulations, high-leverage terminal-phase deviations materially biased structural parameters while remaining insufficiently “extreme” in CWRES space (often < 4), because the non-robust fit partially compensated by inflating the residual variance, thereby shrinking standardized residuals and obscuring the very contamination the diagnostic is intended to detect. Because CWRES are computed with respect to the fitted working residual-error model (here, Gaussian) and its estimated variance, they can become uninformative when the non-robust fit accommodates contamination by inflating σ and shifting structural parameters. In that regime, standardized residuals may shrink even when the raw deviation is large, therefore the absence of large CWRES is not evidence of compatibility with Gaussian errors; it may simply indicate that the model has already shifted parameters and variability to accommodate contaminated points. Collectively, these findings argue against CWRES thresholding as the primary mechanism for outlier management and support likelihood-based robust residual modeling as the default strategy for routine PopPK inference.

This study was motivated by a practical gap in routine pharmacometric workflows: influential outliers are common, yet the dominant operational response in many analyses remains residual screening rather than explicit robust likelihoods. Although Student’s t residual models are widely regarded as an effective solution, ^2, 18, 20, 21^ their use in applied pharmacometrics appears limited in practice, in part because implementation is perceived as burdensome, particularly in tools or workflows requiring custom likelihood specification and in simulation- or sampling-based Bayesian settings where computational efficiency becomes salient. This creates a strong incentive to seek simpler heavy-tailed substitutes. The Laplace and generalized error (GED/exponential-power) families are attractive candidates because they are heavier-tailed than Gaussian errors and admit simple closed-form likelihoods, making them straightforward to implement and potentially more computationally efficient. ^23, 25^ Our results show that this simplicity–robustness trade-off is real: exponential-tail alternatives can reduce sensitivity relative to Normal errors under some contamination patterns, but they do not consistently match Student’s t when deviations are highly influential, particularly when outliers occur at high-leverage time points such as the terminal phase, highlighting the need to distinguish when exponential-tail models are adequate versus when power-law tails are required.

Figure 2 supports the initial plausibility of Laplace and GED as practical alternatives. Relative to the Normal distribution, both families have heavier tails and thus relax the quadratic penalization of Gaussian residual errors. By replacing squared-error loss with exponential-tail decay, these models down-weight moderate residual excursions more gently, behaving qualitatively similarly to Student’s t when deviations are not extreme. Consistent with this expectation, our clean-data simulations showed that all four likelihood specifications (Normal, Student’s t, Laplace, and GED) achieved comparable parameter recovery in the absence of outlier contamination. These observations supported our working hypothesis that Laplace or GED could provide a useful robustness gain with substantially simpler likelihood expressions in custom-likelihood environments.

However, the contaminated simulations and the real-world caffeine drug-drug interaction case study demonstrate that robustness is regime-dependent. Laplace and GED offered meaningful improvement over Gaussian errors under mild-to-moderate contamination, but they did not consistently match Student’s t when deviations became severe, especially when outliers occurred at high-leverage time points such as the terminal phase. This separation is mechanistically consistent with tail behavior. Laplace and GED remain exponential-tail likelihoods, and even when heavier than Gaussian, their tails still decay exponentially; for gross residuals, they can therefore remain overly punitive, reintroducing pressure for the estimation algorithm to accommodate contaminated observations through compensatory model adjustments. In contrast, Student’s t achieves robustness through power-law tails, assigning materially higher probability to extreme deviations and thereby reducing the incentive to absorb outliers through structural distortion.

When the assumed residual model assigns near-zero probability to an extreme observation, mixed-effects estimation commonly responds through compensatory adjustments that erode interpretability. ^26^ One failure mode is structural parameter drift, where fixed effects (e.g., clearance and volume) shift to reduce the residual associated with the contaminated point, biasing clinically meaningful quantities. A second is variance inflation, where residual error and/or IIV increase to “absorb” extreme deviations, often yielding physiologically implausible subject-level estimates while simultaneously masking contamination in standardized residual diagnostics. ^27^ Both behaviors emerged as contamination severity increased in our simulations, and the caffeine case study provides a concrete illustration: terminal-phase spikes distorted the fitted elimination slope when the residual model could not adequately tolerate those deviations. By maintaining non-negligible likelihood for extreme residuals, Student’s t reduces the need for these compensatory mechanisms, allowing outlying observations to be down-weighted without reshaping the underlying kinetics.

Taken together, these findings support pragmatic guidance that is compatible with real-world constraints while remaining statistically principled. When deviations are expected to be mild (e.g., assay noise or occasional moderate measurement errors), Laplace or GED may provide acceptable robustness improvements over Gaussian residuals and may be easier to implement in constrained environments. However, when influential outliers are plausible, Student’s t should be treated as the default robust residual model to preserve parameter interpretability and control bias. A key operational advantage of Student’s t is its adaptivity: the estimated degrees of freedom (ν) provides an internal, data-driven indicator of tail heaviness, large ν implies near-Gaussian behavior, whereas low ν indicates meaningful heavy-tailed contamination. This “adaptive robustness” is difficult to replicate with fixed-form exponential-tail alternatives without introducing additional complexity, and it provides a practical justification for treating Student’s t as the primary likelihood-based tool for outlier-robust PopPK inference.

## Conclusion

This study systematically benchmarked four residual likelihood specifications for outlier handling in population PK modeling (Normal, Laplace, GED/exponential-power, and Student’s t). Two conclusions emerge. First, residual-based screening approaches such as CWRES thresholding are not a reliable safeguard for identifying or mitigating influential PK outliers, particularly when deviations occur at high-leverage time points and the model can partially mask contamination through variance inflation and parameter drift. Accordingly, robust pharmacometric analyses should not rely on CWRES cutoffs as the primary outlier-handling strategy. Second, among the likelihood families evaluated, Student’s t residual errors provide the most consistent robustness, attributable to their power-law tail behavior and the adaptive degrees-of-freedom parameter that allows the model to revert toward Gaussian behavior when contamination is minimal. Collectively, these results support a practical shift in routine pharmacometric workflows: replacing CWRES-driven outlier handling with Student’s t-based residual modeling as the default approach for robust PopPK inference when outlier contamination is plausible.

### Consent for publication

All the authors have reviewed and concurred with the manuscript.

## Authors’ contributions

Y.C. and Y.L. contributed to conception and design; Y.C. and Y.L. contributed to acquisition of data; Y.C. and Y.L. contributed to analysis; all authors contributed to interpretation of data; Y.C. and Y.L. drafted and revised the article. Both authors made substantial contributions to conception and design, acquisition of data, or analysis and interpretation of data; took part in drafting the article or revising it critically for important intellectual content; agreed to submit to the current journal; gave final approval of the version to be published; and agreed to be accountable for all aspects of the work.

---

**Supplementary Figure S1.**
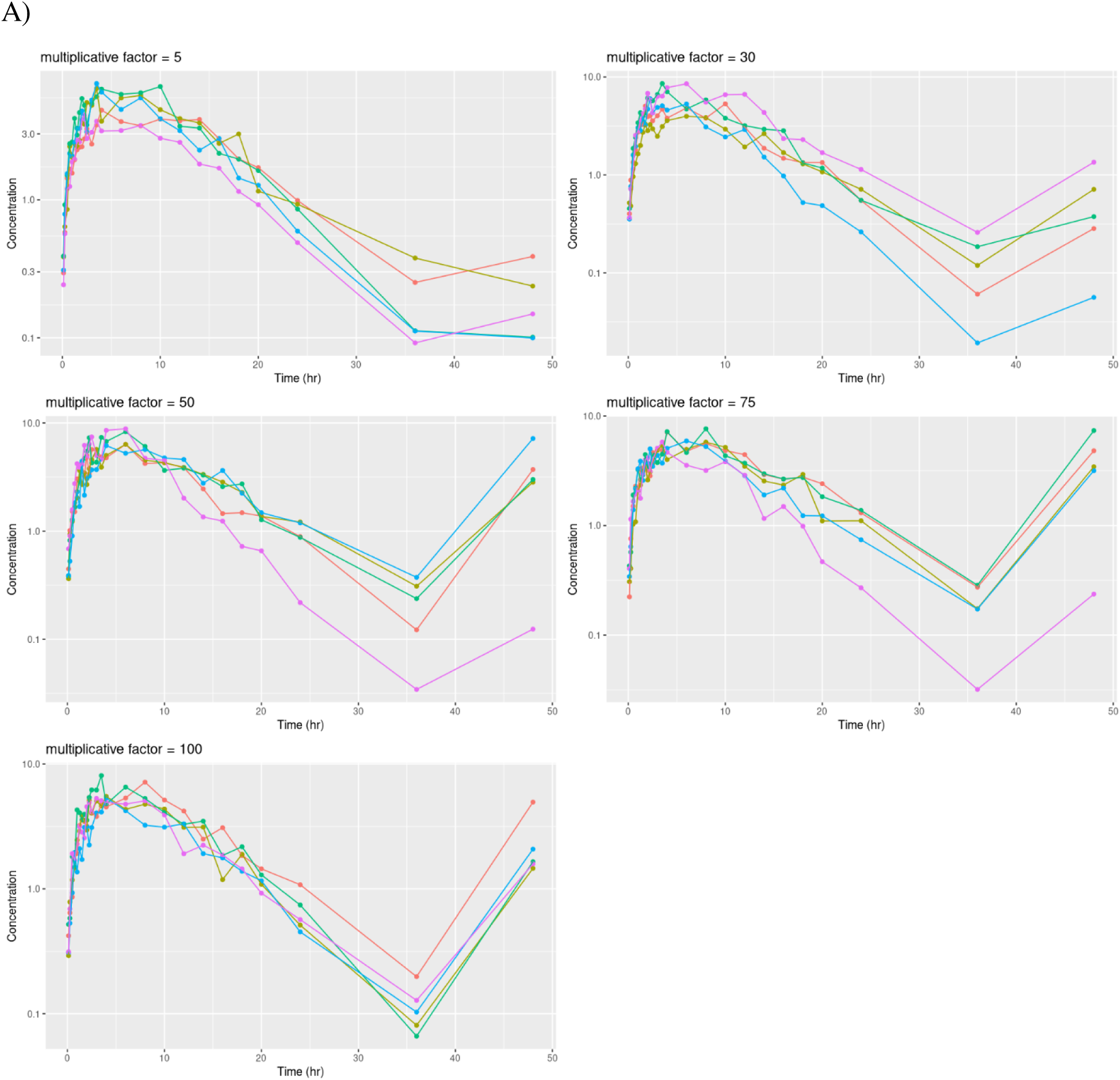
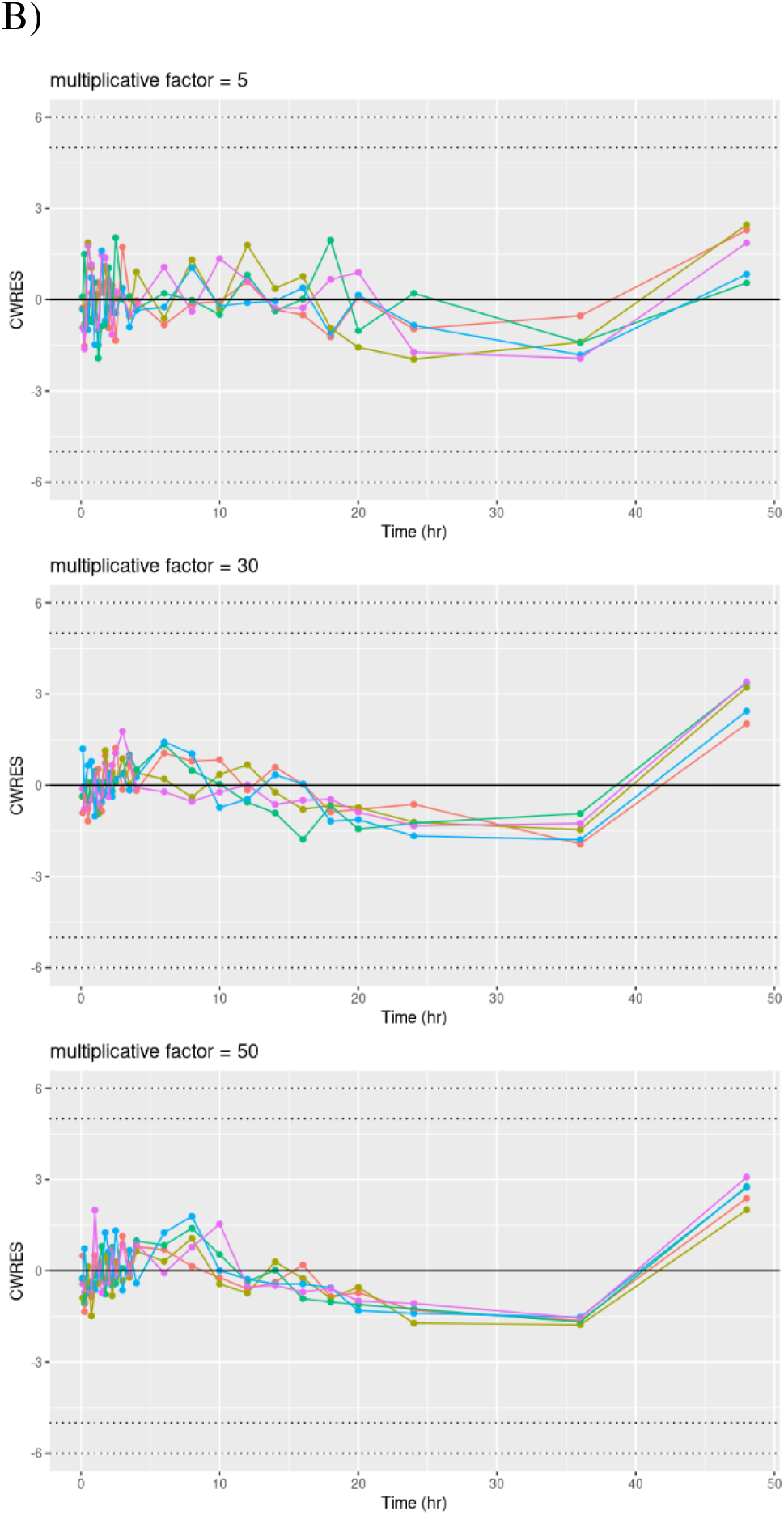
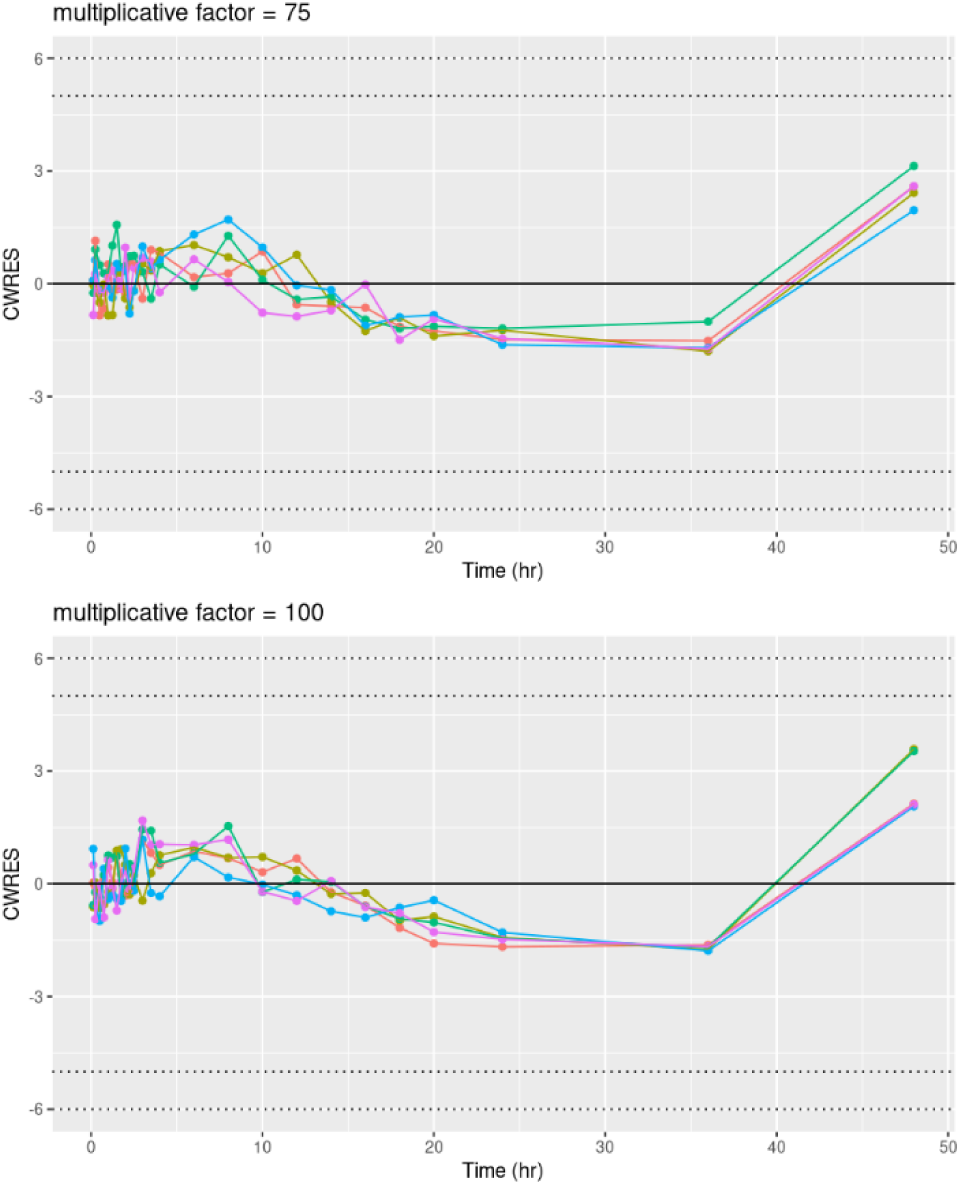
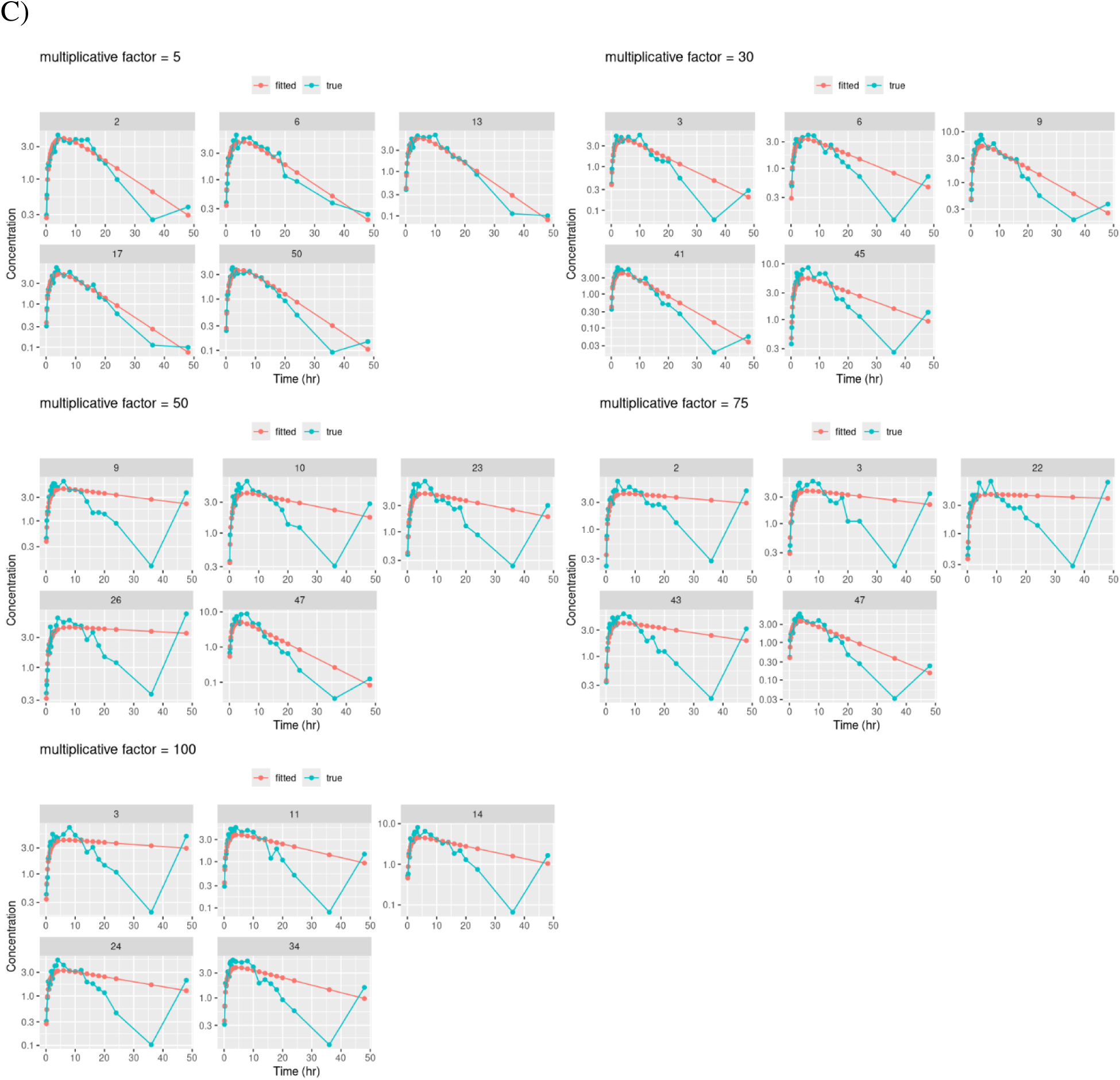
Impact of influential terminal-phase outliers on CWRES diagnostics and pharmacokinetic modeling across multiplicative inflation factors (5, 30, 50, 75, and 100). (A) Simulated concentration–time profiles for five representative subjects. (B) Corresponding CWRES time courses for the same subjects. (C) Observed concentration–time profiles (green) overlaid with model-predicted profiles (red) for the same subjects. Dotted horizontal lines denote the |CWRES| thresholds of 5 and 6.

**Supplementary Figure S2.**
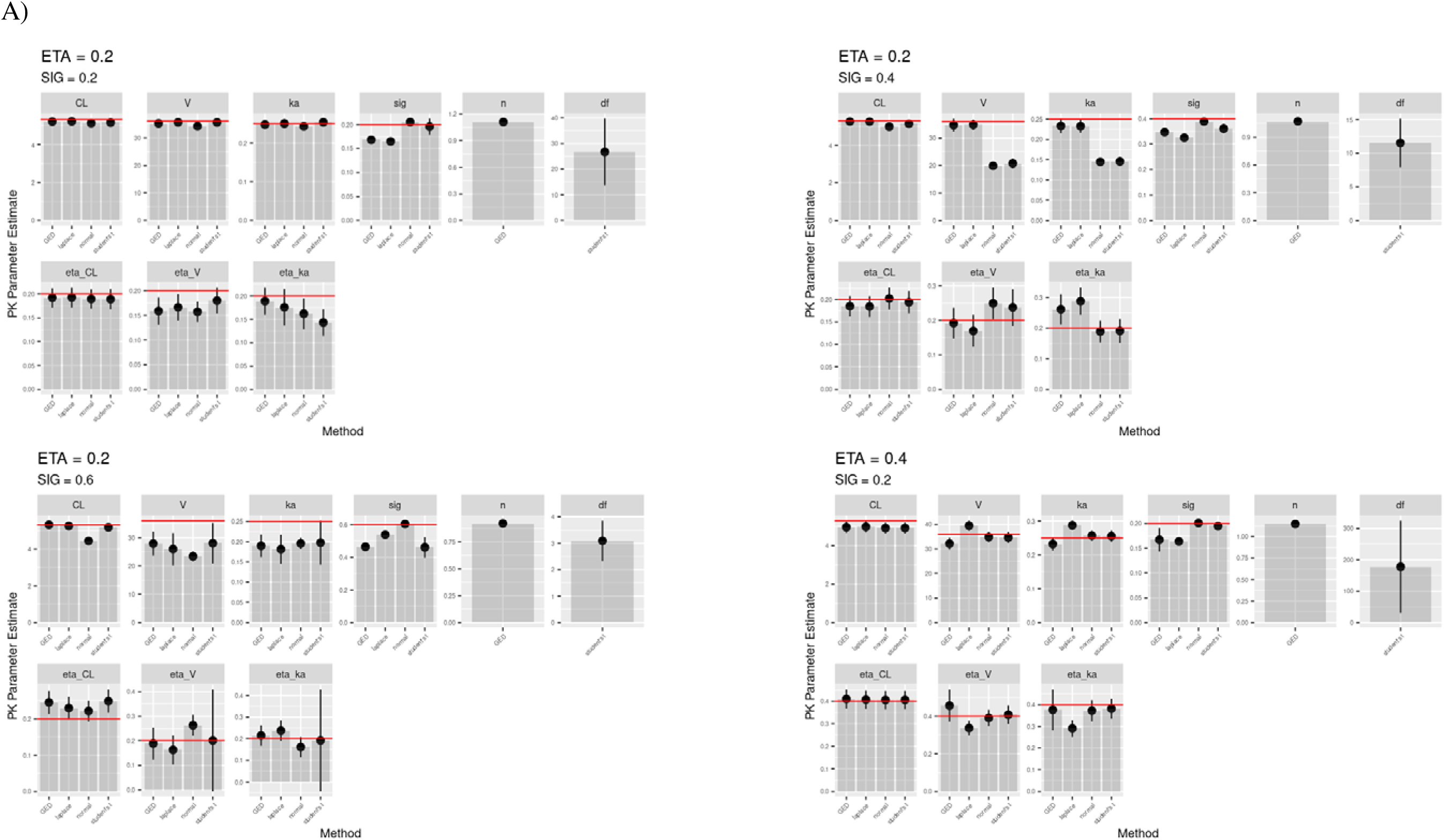
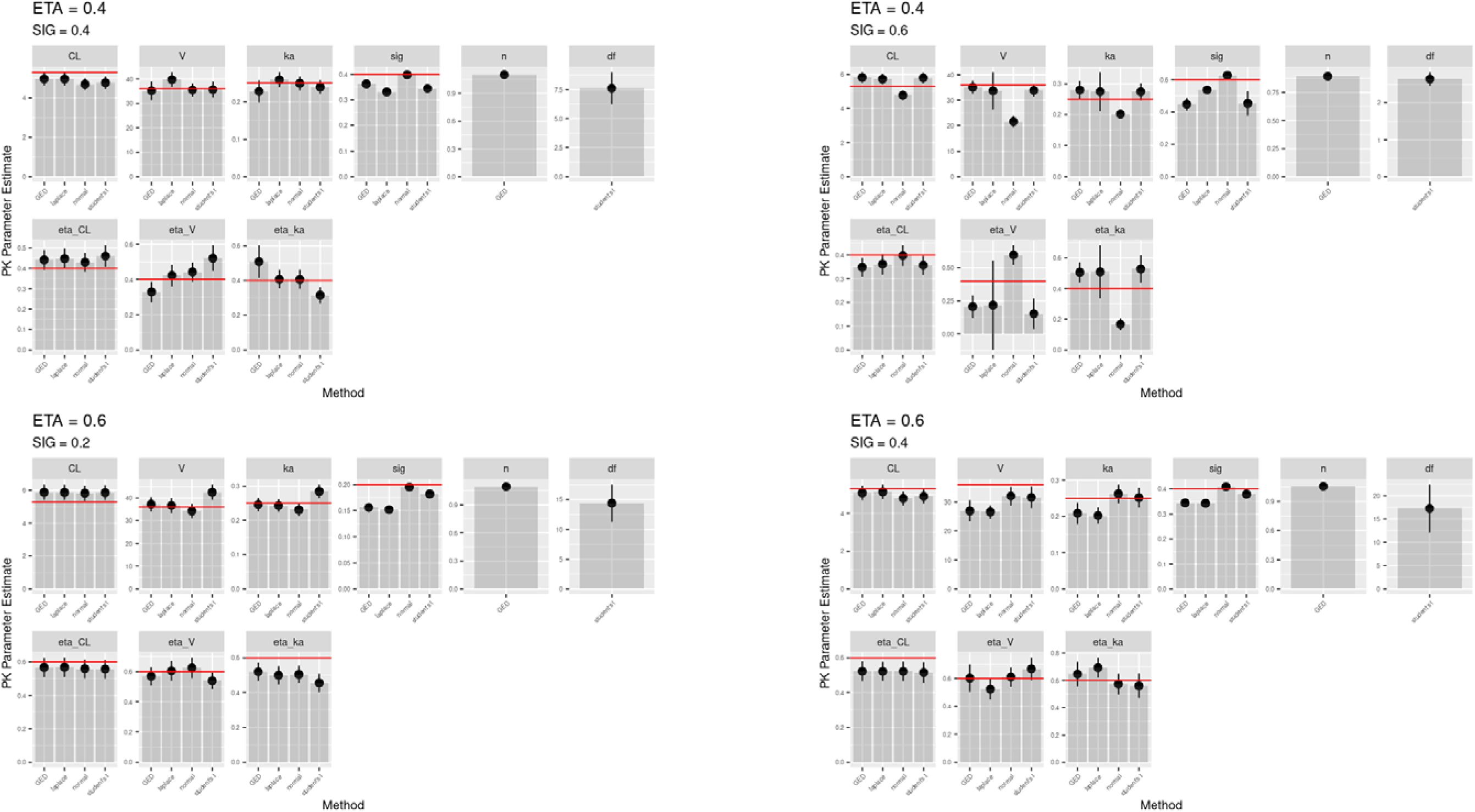
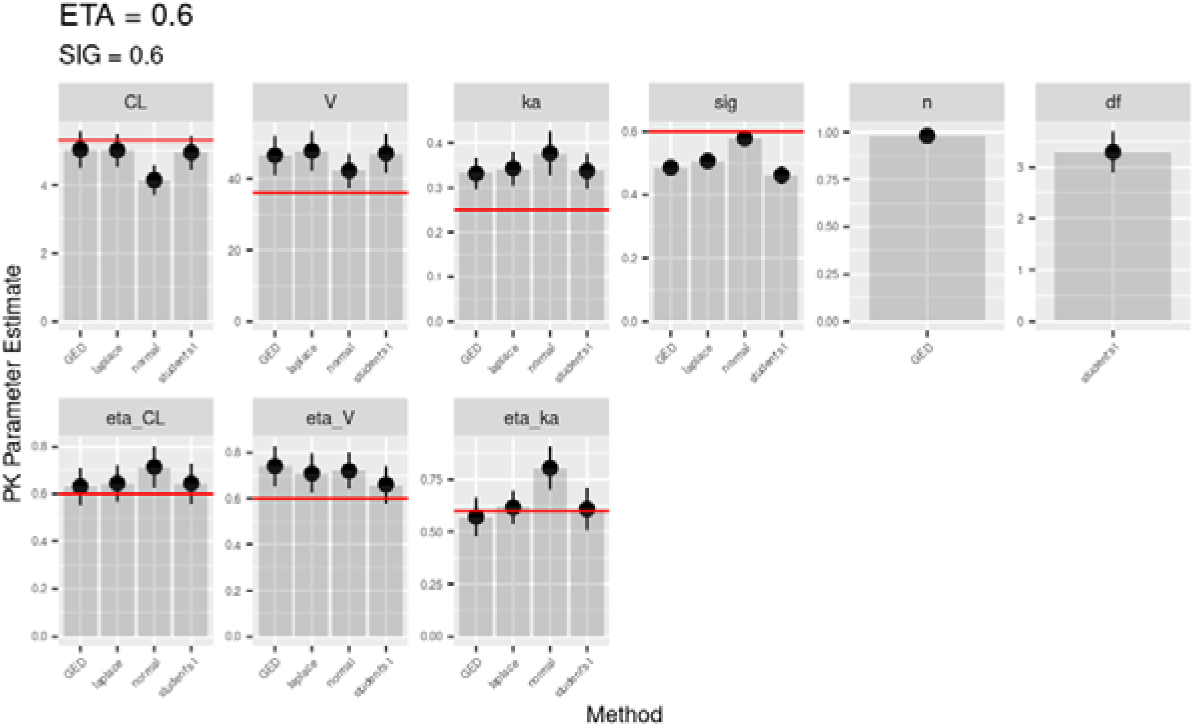
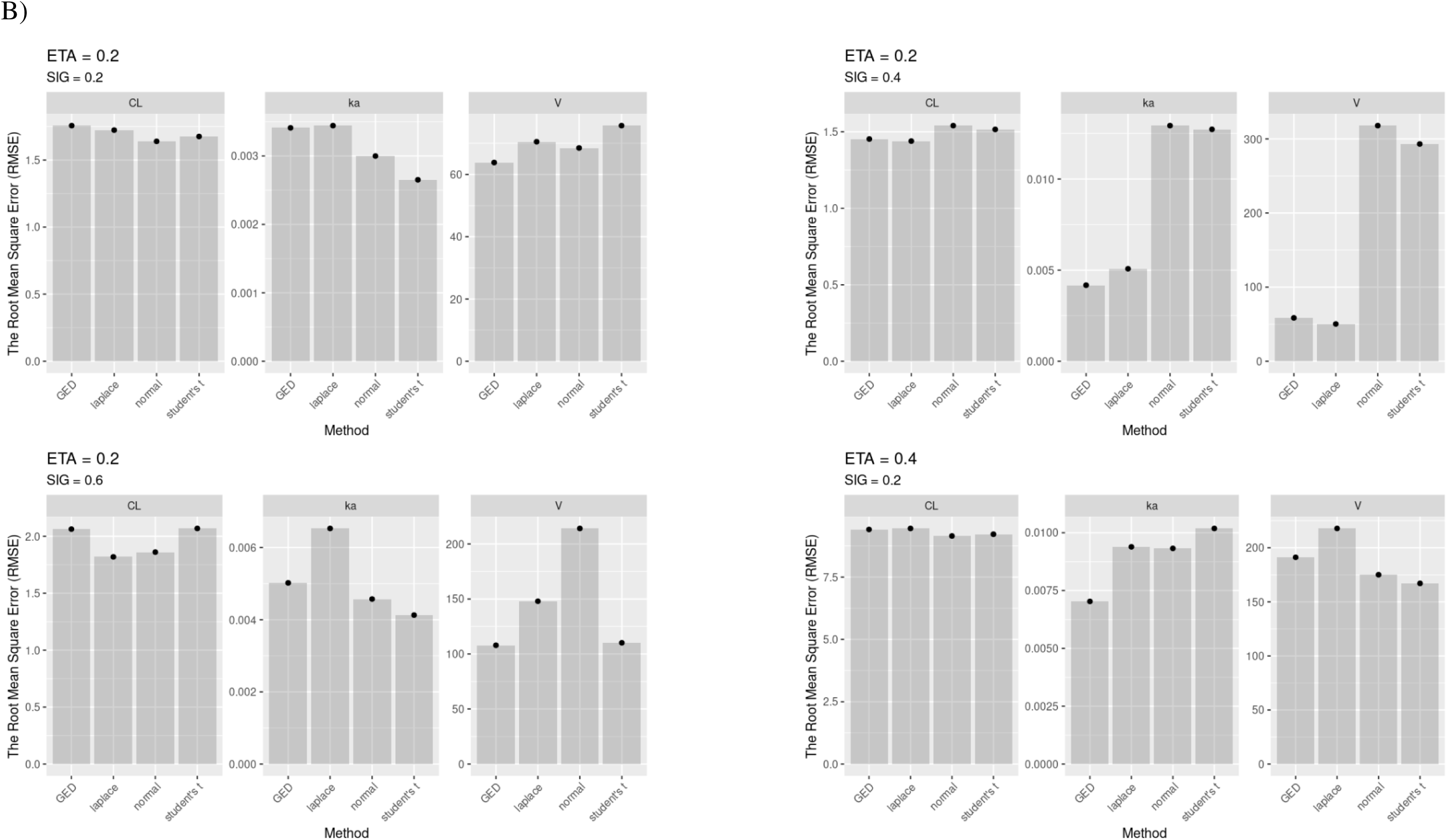
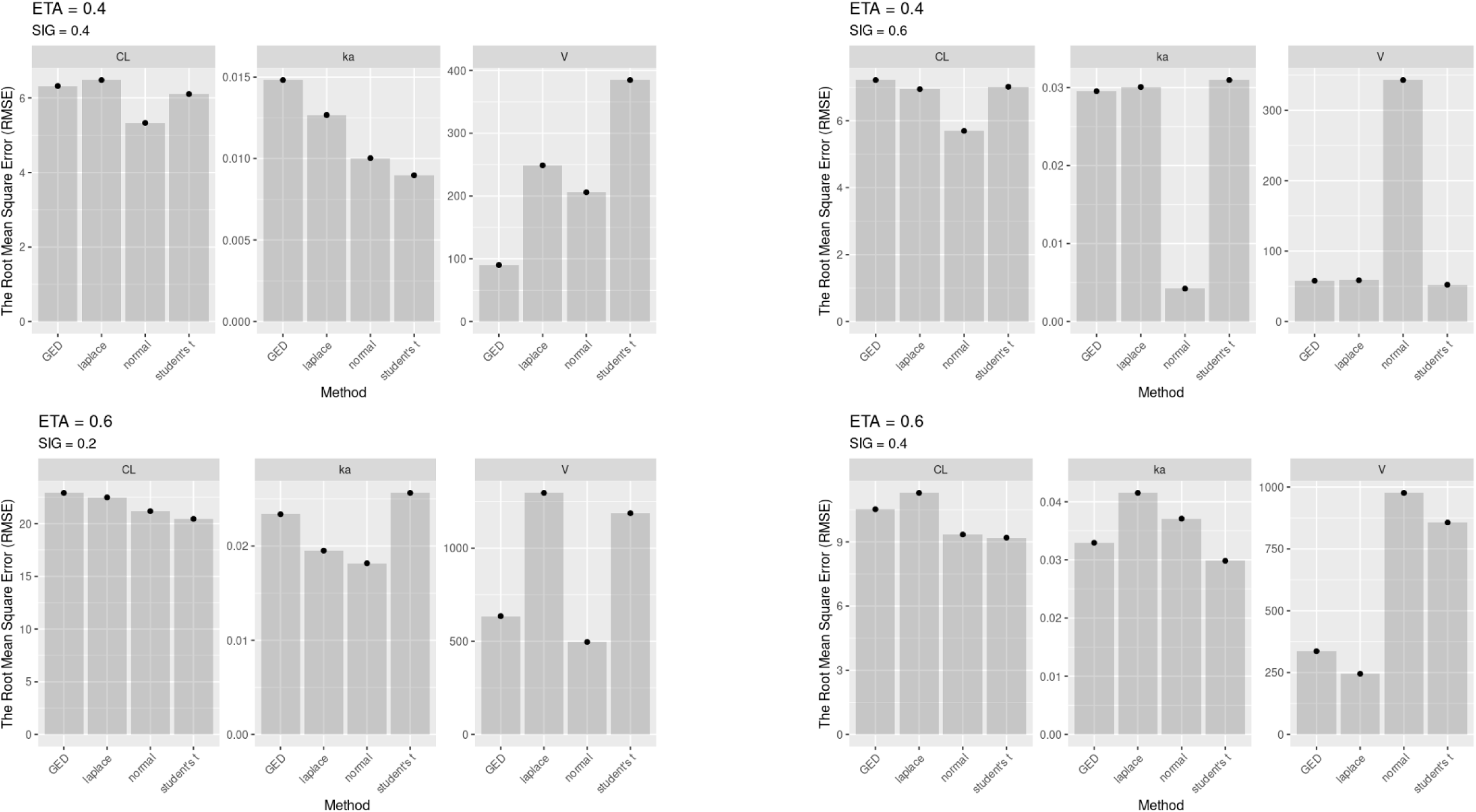
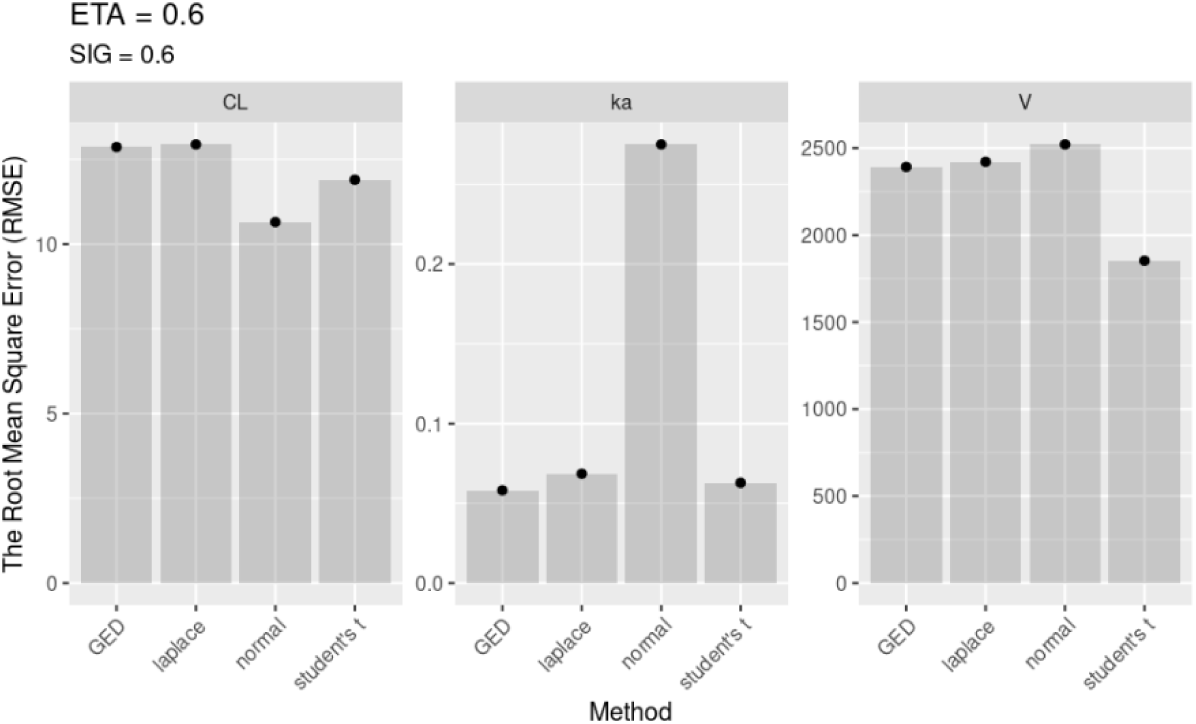
Baseline simulation without outliers: population- and individual-level PK parameter recovery under Normal, Student’s t, Laplace, and GED residual-error models across a wider range of variability settings. Proportional inter-individual variability (IIV) and proportional residual error were varied in combination (e.g., 0.2, 0.4, and up to 0.6).

**Supplementary Figure S3.**
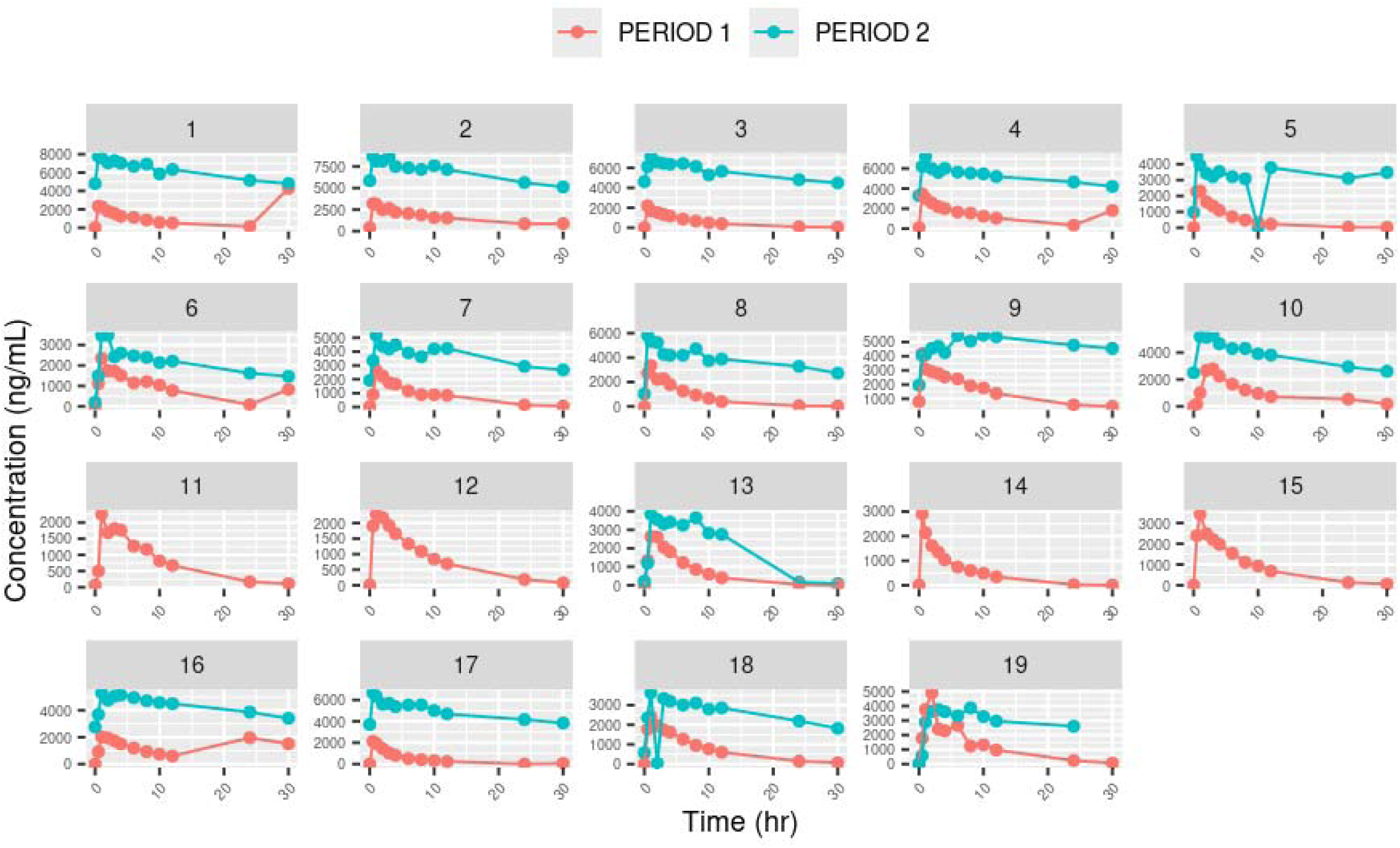
Individual caffeine concentration–time profiles from the drug–drug interaction case study by period (Period 1: absence; Period 2: presence of perpetrator).

